# Reproducible and fully automated testing of nocifensive behavior in mice

**DOI:** 10.1101/2023.04.13.536768

**Authors:** Christopher Dedek, Mehdi A. Azadgoleh, Steven A. Prescott

## Abstract

Pain in rodents is often inferred from their withdrawal to noxious stimulation, using the threshold stimulus intensity or response latency to quantify pain sensitivity. This usually involves applying stimuli by hand and measuring responses by eye, which limits reproducibility and throughput to the detriment of preclinical pain research. Here, we describe a device that standardizes and automates pain testing by providing computer-controlled aiming, stimulation, and response measurement. Optogenetic and thermal stimuli are applied to the hind paw using blue and infrared light, respectively. Red light delivered through the same light path assists with aiming, and changes in its reflectance off the paw are used to measure paw withdrawal latency with millisecond precision at a fraction of the cost and data processing associated with high-speed video. Using standard video, aiming was automated by training a neural network to recognize the paws and move the stimulator using motorized linear actuators. Real-time data processing allows for closed-loop control of stimulus initiation and termination. We show that stimuli delivered with this device are significantly less variable than hand-delivered stimuli, and that reducing stimulus variability is crucial for resolving stimulus-dependent variations in withdrawal. Slower stimulus waveforms whose stable delivery is made possible with this device reveal details not evident with typical photostimulus pulses. Moreover, the substage video reveals a wealth of “spontaneous” behaviors occurring before and after stimulation that can considered alongside withdrawal metrics to better assess the pain experience. Automation allows comprehensive testing to be standardized and carried out efficiently.

## INTRODUCTION

Measuring withdrawal from noxious stimuli in laboratory rodents is a mainstay of preclinical pain research (Barrot, 2012; Deuis et al., 2017; Gregory et al., 2013; Le Bars et al., 2001). Testing is often conducted on the hind paw, in part because many chronic pain models are designed to increase paw sensitivity through manipulations of the paw or the nerves innervating it (Abboud et al., 2021; Burma et al., 2017; Gregory et al., 2013; Jaggi et al., 2011). Measuring evoked pain with withdrawal reflexes has been criticized (Mogil & Crager, 2004) since ongoing (non-evoked) pain is a bigger clinical problem (Backonja & Stacey, 2004), but tactile and thermal sensitivity are altered in many chronic pain conditions (Maier et al., 2010) and allodynia and spontaneous pain intensity tend to be correlated in human studies (Koltzenburg et al., 1994; Rowbotham & Fields, 1996) and in some (Pitzer et al., 2016) but not all (Mogil et al., 2010) mouse studies. Furthermore, sensory profiling is useful for stratifying patients in clinical trials (Baron et al., 2022; Edwards et al., 2016) and altered sensitivity is central to diagnosing certain conditions, e.g. fibromyalgia (Arnold et al., 2019). It logically follows that ongoing pain should be assessed in addition to, not instead of, evoked pain (Negus, 2019). Doing so would provide a more complete picture, including the relationship between evoked and ongoing pain. We must also identify and correct the most problematic aspects of this testing; for instance, outcomes of the hot water tail flick test was shown to depend more on who conducts the testing than on any other factor (Chesler et al., 2002). This likely generalizes to other behavioral tests but has received scant attention compared with other factors like sex (Sadler et al., 2021). Outdated technology and poorly standardized testing protocols contribute to the oft-cited reproducibility crisis (Mogil, 2017) and are long overdue for transformative improvements.

Preclinical pain tests typically measure withdrawal threshold using brief repeated (incrementing) stimuli like von Frey filaments, or sustained stimuli like radiant heat. The stimulus intensity (force or skin temperature) at which withdrawal occurs is assumed to be the lowest intensity perceived as painful (i.e. pain threshold) notwithstanding certain caveats (Le Bars et al., 2009). Of course, withdrawal might not always be triggered by pain, and focusing on threshold fails to consider pain intensity over a broader range of stimulus intensities. Recent studies have quantified responses to suprathreshold mechanical stimulation using high-speed video (Abdus-Saboor et al., 2019; Fried et al., 2020; Jones et al., 2020), but despite precise response measurement, stimuli were delivered by hand and throughput was low. Resolving subtle changes in pain sensitivity requires that stimulus-response relationships be measured with high resolution (which requires reproducible stimulation and precise response measurement), over a broad dynamic range, and with reasonable efficiency (throughput). Improvements in one factor (stimulation, response measurement, throughput, etc.) may come at the expense of other factors. The best compromise depends on the particular experiment, but improving reproducibility *and* throughput would be a huge benefit.

Optogenetics has provided an unprecedented opportunity to study somatosensory coding, including nociception. Expressing actuators like channelrhodopsin-2 (ChR2) in genetically defined subsets of afferents allows those afferents to be selectively activated or inhibited with light applied through the skin (transcutaneously) or directly to the nerve or spinal cord using more invasive methods (Copits et al., 2016; Xie et al., 2018). Afferents can be optogenetically activated with patterns not possible with somatosensory stimulation; for instance, mechanical stimuli that activate Aδ high-threshold mechanoreceptors (HTMRs) normally also activate low-threshold mechanoreceptors (LTMRs), so it was only by expressing channelrhodopsin-2 (ChR2) selectively in HTMRs that HTMRs can be activated in isolation (Arcourt et al., 2017). Causal relationships between afferent activation patterns and perception/behavior can be thoroughly tested in this way. Elucidating those relationships is key to understanding physiological pain and how pathology disrupts normal coding, facilitating development of targeted therapies. Optogenetics has been used for basic pain research but, despite its potential, has not yet been adopted for drug testing (Woolf, 2020). Transcutaneous photostimulation is amenable to high-throughput testing but, like tactile and thermal stimuli, is hard to apply reproducibly in behaving animals.

We sought to improve the reproducibility of transcutaneous photostimulation while also streamlining response measurement in order to increase throughput. We developed a device able to deliver photostimuli consistently and measure withdrawal latency automatically with millisecond precision. Using this device, we show that withdrawal latency correlates inversely with the intensity of photostimulus pulses, and that photostimulus ramps reveal differences not seen with pulses. We also incorporated an infrared laser for radiant heating, allowing the same device to measure optogenetic and thermal sensitivity. With a clear view of the mouse from below, a neural network was trained to recognize the paw and aim the stimulator, thus fully automating the testing process. The substage video also provides a wealth of data about spontaneous behaviors and their potential association with evoked pain.

## RESULTS

Mice are kept individually in enclosures on a clear platform with the stimulator underneath (**Fig. 1A**). Different enclosures were tested including a novel design in which the mouse is transferred from its home cage in a clear plexiglass tube, which is then turned vertically and slid into an opaque ceilinged cubicle for testing (**Fig. 1B**). Tube handling is less stressful than other handling methods (Gouveia & Hurst, 2013, 2019; Hurst & West, 2010) and using separate tubes facilitates addition/removal of individual mice from the multi-mouse cubicle. For high-speed video, which required the mouse to face left to view the stimulated paw in profile, we used a narrow rectangular chamber with clear walls on the front and left side.

**Figure 1.**
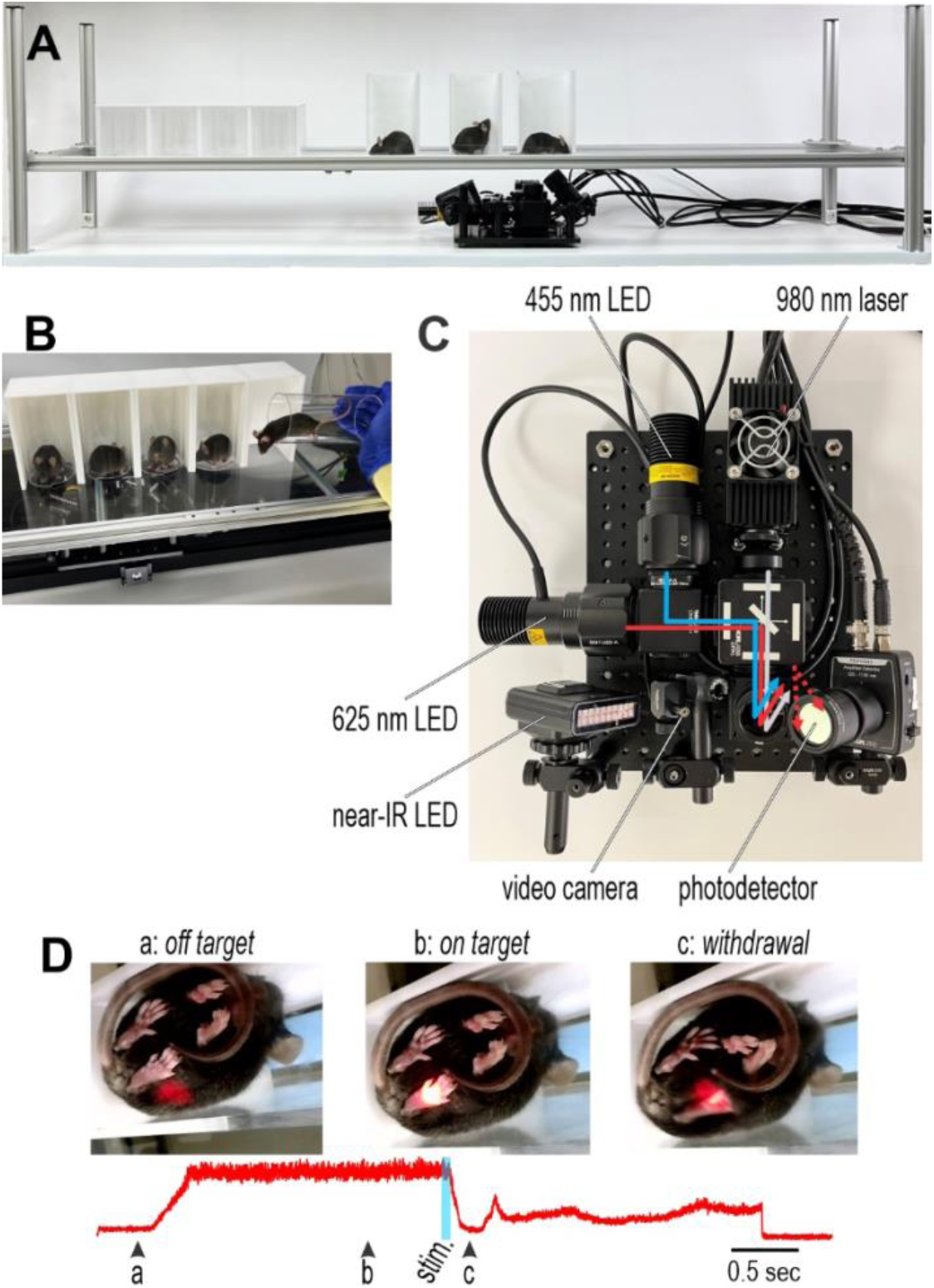
Overview of equipment. (**A**) Mice are kept separately in enclosures on a plexiglass platform. The stimulator remains at a fixed distance below the platform and is positioned by hand (version shown here) or by motorized actuators (see Fig. 6). **(B)** Novel enclosure design. Mice are transferred from their home cage to the platform in a clear plexiglass tube, which is placed vertically on the platform and slid back into an opaque cubicle. **(C)** Top view of photostimulator. Blue and IR light is used for optogenetic stimulation and radiant heating, respectively. Red light is used to help aim and to detect paw withdrawal. All wavelengths are combined into a single beam delivered to the same spot, but their intensities are independently controlled by computer. See **Supplementary Figure 1** for details. **(D)** Sample frames from substage video (**Video 1**) before the stimulator is properly aimed (a), after aiming (b), and after paw withdrawal (c). Red trace shows the intensity of reflected red light measured by the photodetector.

**Figure 1C** shows the stimulator viewed from above. Blue light for optogenetic activation using ChR2, IR light for thermal stimulation (radiant heating), and red light for aiming and response measurement are combined into a single beam using dichroic mirrors (**Supplementary Fig. 1**). The beam is directed vertically and focused to a spot 5 mm in diameter on the platform above. An adjacent borescope collects video from below (substage) while a photodetector measures red light reflected off the paw (**Fig. 1C**). A near-IR LED helps improve lighting during high-speed video, which tends to be dark because of the short exposure time. The stimulator is translated manually or by motorized actuators (see Fig. 6) using the substage video to aim.

Since all wavelengths converge on the same spot, red light is turned on prior to initiating photostimulation (with blue or IR light) to verify where photostimuli will hit, thus providing visual feedback to optimize aiming (**Fig. 1D**; **Video 1**). Rodents are typically assumed not to see red light (De Farias Rocha et al., 2016); though some evidence contradicts this (Nikbakht & Diamond, 2021; Niklaus et al., 2020), we never observed any behavioral response to red light (little of which likely reaches the eyes when applied to the hind paw), suggesting that the aiming phase does not provide mice any visual cue about the forthcoming photostimulus. Reflectance of red light off the paw is measured by the adjacent photodetector (red trace). Maximization of the reflectance signal can be used to optimize aiming (compare frames a and b). This reflectance signal is stable while the paw and stimulator are immobile but changes when the paw is withdrawn (frame c), thus enabling measurement of withdrawal latency (see Fig. 3). Though too slow to accurately measure fast withdrawals, standard video provides a visual record to rule out gross errors in reflectance-based latency measurements and enables assessment of slower behaviors like guarding and licking (see Fig. 5).

### Reproducible Stimulation

Unaccounted for variations in stimulation fundamentally limit the precision with which stimulus-response relationships can be characterized. LEDs and lasers offer stable light sources but the amount of light hitting a target can vary over time or across trials depending on the accuracy and precision of aiming. When applying light by hand-held fiber optic (as typically done for transcutaneous stimulation), stability of the tester and differences in aiming technique across testers are important. To gauge the importance of aiming, we measured how the amount of light hitting a target depended on the fiber optic’s positioning in the *x*-*y* plane and its distance (*z*) below the platform. Light was delivered through a paw-shaped cut-out to a photodetector facing downward on the platform (to simulate stimulation of a mouse paw; **Video 2**) while controlling fiber optic position with linear actuators. **Figure 2A** shows that light delivery is sensitive to positioning in all three axes, especially in *z* (because light rays diverge from the fiber optic tip).

**Figure 2.**
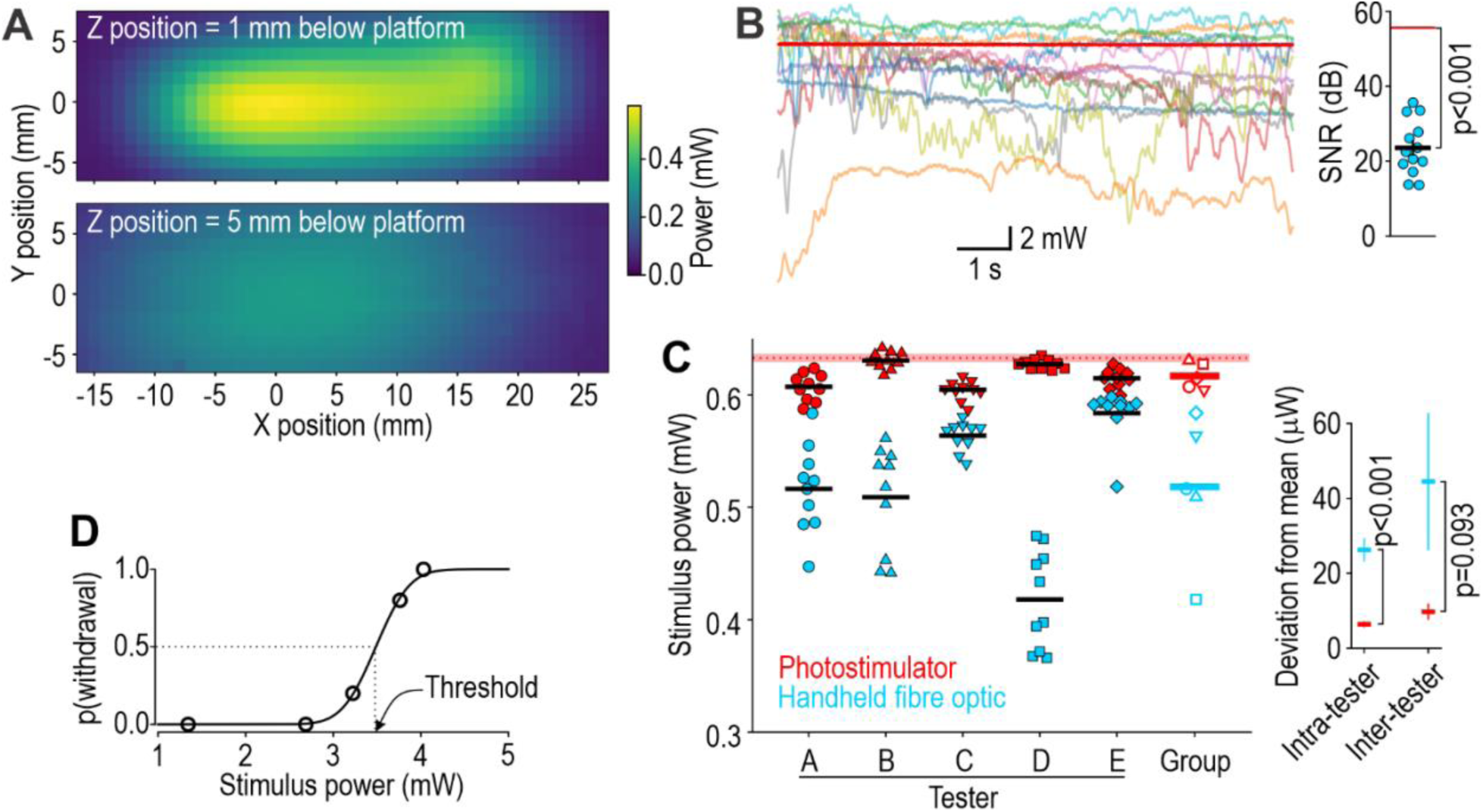
Reproducible photostimulation. For A-C, on-target light was measured by stimulating a photodetector facing down on the platform with a paw-shaped cut-out over its surface (**Video 2**). **(A)** Importance of fiber optic positioning. Fiber optic was mounted on linear actuators to control *x-y* positioning; *x*-axis aligns with long axis of the paw with 0 position centered on the maximal response. Measurements were repeated for two distances (*z*) below the platform. **(B)** Light delivery during 10 s-long stimulus. Signal-to-noise ratio (SNR = mean^2^/variance) when using the stimulator (55.6 dB; red trace) was significantly higher than for the handheld fiber optic (23.5 ± 2.0 dB, mean ± SEM; pale traces, 1 for each of 13 testers) (*T*_12_ = 15.8, *p* < 0.001, one sample *t*-test). **(C)** Light delivery during 100 ms-long pulses. Filled symbols show individual trials with a handheld fiber optic (blue) or stimulator (red); black lines represent intra-tester averages. Average trial-to-trial deviation from each tester’s average was significantly larger for handheld fiber optic (26.3 ± 3.0 mW; mean ± SEM) than for stimulator (6.4 ± 0.7 mW) (*T*_98_ = 6.42, *p* < 0.001, unpaired *t*-test). Open symbols represent intra-tester averages; colored lines represent group average. Tester-to-tester deviation from the group average was larger for handheld fiber optic (44.5 ± 18.2 mW) than for stimulator (9.7 ± 2.0 mW) (*T*_8_ = 1.94, *p* = 0.093). Red line and shading show average light intensity ± SD across 10 trials without moving the stimulator. **(D)** Example input-output curve from one mouse. Five 100 ms-long blue pulses were delivered at each of 5 intensities. Threshold is intensity at 50% probability of withdrawal, as inferred from fitted curve.

To explore the practical consequences of this, we measured light delivery while 13 testers applied a 10 s-long photostimulus by handheld fiber optic (**Fig. 2B**, pale traces). The signal-to-noise ratio (SNR = mean^2^/SD^2^) of 23.5 ± 2.0 dB (group mean ± SEM) was significantly less than the 55.6 dB obtained with the stimulator (red trace) (*T*_12_ = 15.8, *p* < 0.001, one sample *t*-test). The mean stimulus intensity also differed across testers with an inter-tester coefficient of variation (CV = SD/mean) of 18.8%, which is even larger than the average intra-tester CV of 8.9%. In other words, during a sustained photostimulus, temporal variations in light-on-target arise from each tester’s instability, but this variability is compounded by differences in aiming technique across testers.

The same issues affect short (pulsed) stimuli but manifest as trial-to-trial variations. To measure variability across trials, five testers used a handheld fiber optic or the stimulator to deliver ten 100 ms-long pulses to a photodetector (**Fig. 2C**); each pulse was triggered independently. Trial-to-trial deviation of each tester from their individual mean dropped from 26.3 ± 3.0 mW (mean ± SEM) with the handheld fiber optic to 6.4 ± 0.7 mW with the stimulator (*T*_98_ = 6.42, *p* < 0.001, unpaired *t*-test), which represents a 75.6% reduction in intra-tester variance. Deviation of each tester from the group mean fell from 44.5 ± 18.2 mW with the fiber optic to 9.7 ± 2.0 mW with the stimulator (*T*_8_ = 1.94, *p* = 0.093), which represents a 78.2% reduction in inter-tester variance. In other words, using the stimulator increased reproducibility of stimulation across testers and within each tester. This is because the photostimulator’s distance below the platform is fixed and light rays converge, and because aiming is improved by using video and visual feedback from red light.

Even if stimulation is reproducible, behavior is still variable, especially in response to weak stimuli. Indeed, threshold is defined as the stimulus intensity at which withdrawal occurs on 50% of trials. **Figure 2D** shows determination of optogenetic threshold. Reliable aiming combined with precisely controllable LEDs (whose output can be varied in small increments over a broad range) allows one to measure threshold and characterize the broader stimulus-response relationship, assuming responses can be identified clearly and measured precisely.

### Precise Response Measurement

High-speed video is the gold standard for measuring fast behaviors, but acquiring and analyzing those data is complicated and costly. We sought to replace high-speed video by detecting changes in the amount of red light reflected off the paw (see Fig. 1D) using a low-cost photodetector. To validate our method, response latency was determined from high-speed video for comparison with latency determined from the reflectance signal on the same trial (**Fig. 3A**). The stimulated paw was identified (colored dot in sample frames) using DeepLabCut (Mathis et al., 2018) and paw height was measured from each frame. Latency was determined as the time taken for each signal to cross a threshold. **Figure 3B** shows reflectance-based latency measurements plotted against height-based latency measurements; each point represents one trial from data pooled across 10 mice given 100 ms-long blue pulses with intensities spanning a broad range (same data as in Figure 4). The regression line (green, slope = 1.007) follows the equivalence line (dashed, slope = 1), meaning there is no systematic difference between measurement methods. The error rate is low (<2%) for each method (**Supplementary Fig. 2**). Beyond avoiding an expensive high-speed camera and the challenges of filming the mouse in profile to assess paw height, the reflectance signal can be processed in real-time to enable closed-loop termination of photostimuli. High-speed video is not, therefore, essential for precise latency measurements, and though such video provides additional information (Jones et al., 2020), slower features of the response can be assessed using regular video.

**Figure 3.**
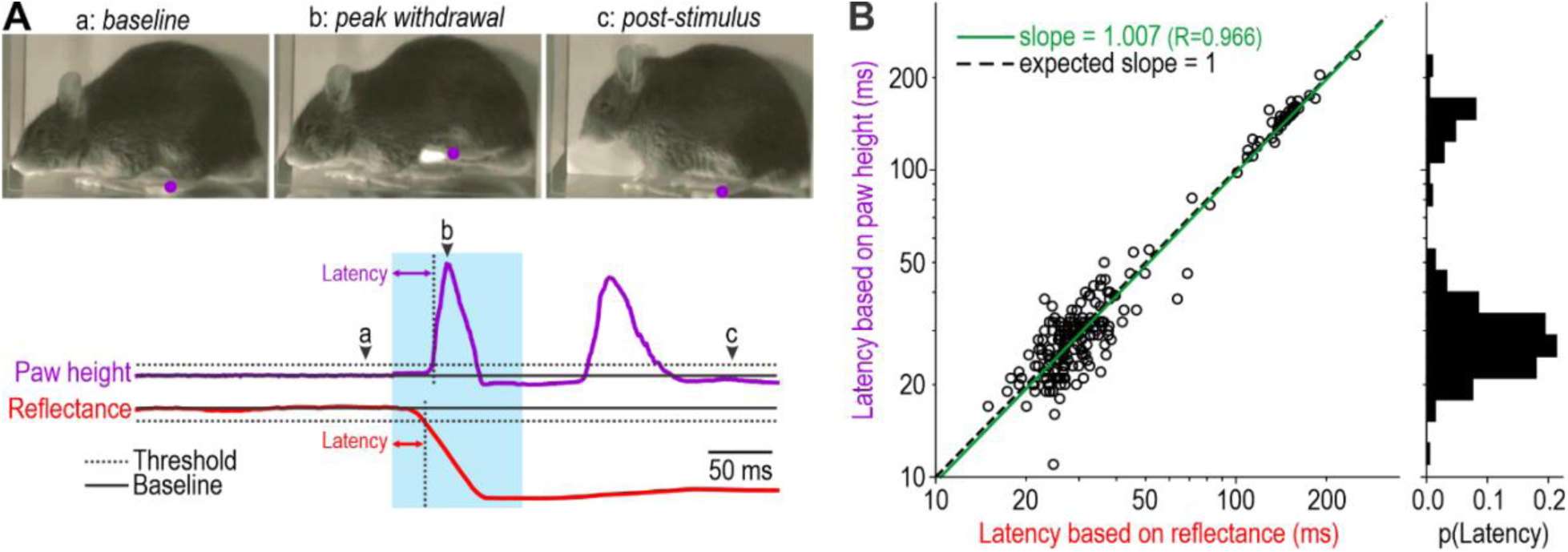
Precise measurement of withdrawal latency. (**A**) Paw withdrawal latency was measured by two methods. Paw height (purple) was extracted from high-speed video (1 kHz) and is plotted alongside intensity of reflected red light (red) measured by the substage photodetector (1 kHz). Sample frames are shown before (a), during (b) and after (c) stimulation with position of stimulated paw (as tracked by DeepLabCut) summarized by a purple dot (**Video 3**). Withdrawal latency was determined as delay from stimulus onset until each signal crossed a threshold (dotted line) defined relative to its baseline (solid line). **(B)** Comparison of latencies measured from each signal. Data points, each representing a single trial, fell along line representing equivalence (dashed, slope = 1), yielding a regression line (green) with slope = 1.007. Data are shown on a log scale to help illustrate their bimodal distribution (histogram on right). Starting from 218 trials from 10 mice, 7 trials were excluded (3.2%) based on errors identified by visual inspection of raw data (see **Supplementary Fig. 2**). Data here are from two experimenters who each tested 5 mice from two genotypes (Advillin-ChR2 and TrpV1-ChR2) of either sex; no effect of sex or genotype was observed, and data were pooled.

**Figure 4.**
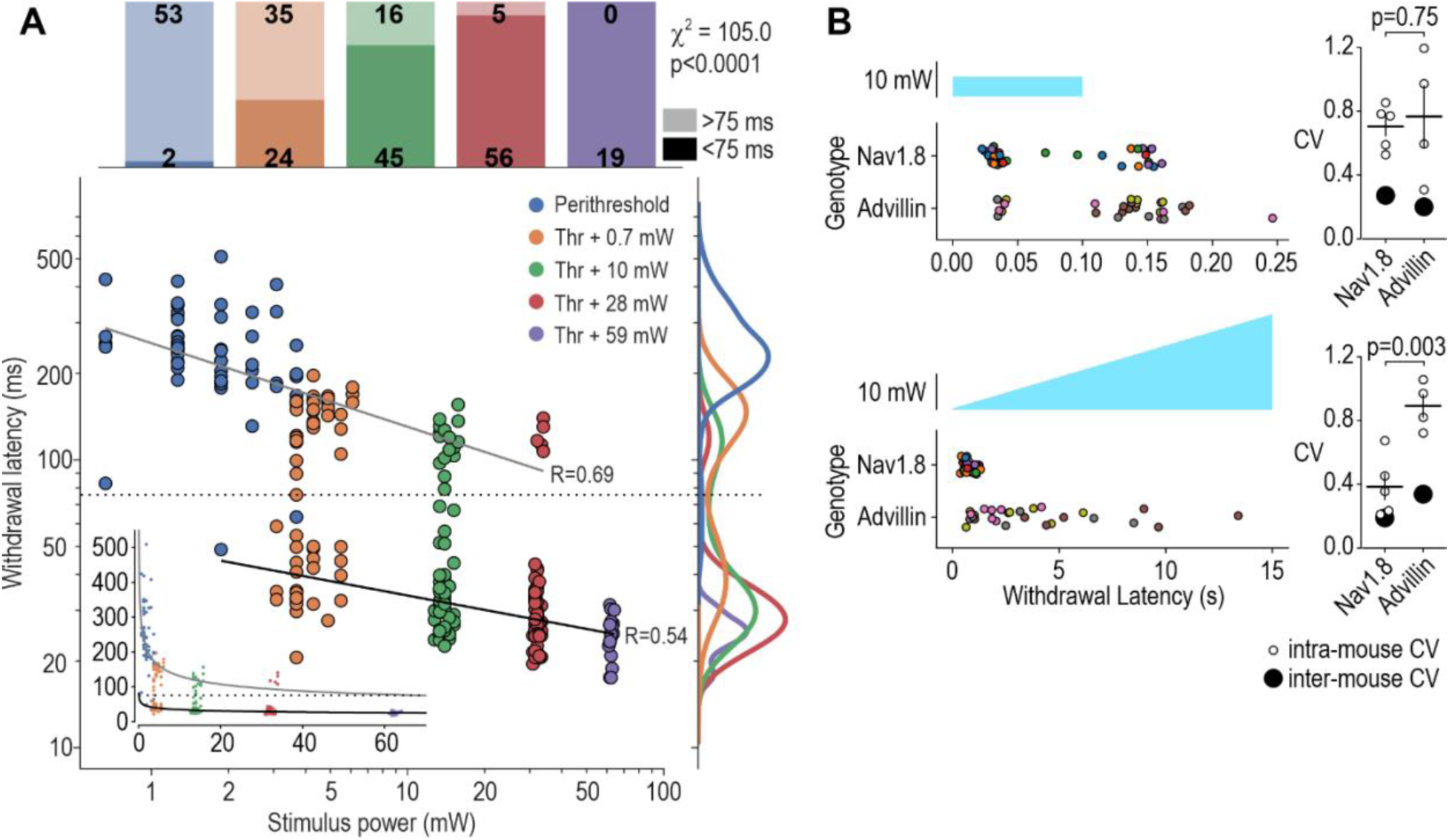
Stimulus-response characterization. (**A**) Impact of photostimulus intensity. Threshold intensity for 100 ms-long blue photostimuli was determined in 10 mice (same as Fig. 3), and then each mouse was re-tested at four intensities relative to its threshold (3 trials per intensity for all but the highest intensity, which was tested only once); withdrawal latency was recorded for all responses. Horizontal variations in blue data points reflect the various intensities used for threshold determination (and are accentuated by the log scale); horizontal variations for other colors reflect inter-animal variability in threshold. Inset shows data on linear scales. Latencies exhibit a bimodal distribution (see histogram on right) with a preponderance of slow (>75 ms) responses at low intensities and fast (<75 ms) responses at high intensities (see stacked bars at top); proportions varied significantly with stimulus intensity (χ^2^ = 105.01, *p* < 0.0001). Black and grey lines show separate linear regressions for fast and slow responses, respectively, and correspond to curves on linear scales (see inset). Fast, *y* = 49.8 *x*^-0.17^. Slow, *y* = 254.8 *x*^-0.29^. **(B)** Impact of photostimulus kinetics. Mice of two genotypes were tested with 100 ms-long pulses (top) and 15 s-long ramps (bottom) (**Video 4**). Data from each mouse are shown in a different color. Pulse-evoked responses exhibited a bimodal latency distribution like in A; genotypes were similar in this regard although distributions differed (*D* = 0.403, *p* = 0.009, two-sample Kolmogorov-Smirnov test). Intra-mouse CV did not differ between genotypes (*T*_7_ = 0.335, *p* = 0.75, unpaired t-test). Ramp-evoked responses occurred with much longer latencies (note difference in timescale) and were significantly more variable in Advillin-ChR2 mice (*D* = 0.714, *p* = 3.67 × 10^-8^). Intra-mouse CV differed significantly between genotypes (*T*_7_ = 4.37, *p* = 0.003).

### Characterizing stimulus-response relationships

Minimizing variability in stimulus delivery and response measurement maximizes discrimination of small biological differences; indeed, an input-output relationship is obscured by poorly controlled input or poorly measured output adding noise respectively to the *x*- and *y*-positions of constituent data points. To explore how well our device reveals stimulus-dependent variations in withdrawal latency, we titrated photostimulus intensity to determine the optogenetic threshold in each of 10 mice (see example in Fig. 2D). Then, using intensities at defined increments relative to each mouse’s threshold, we measured withdrawal latency as a function of photostimulus intensity (**Fig. 4A**). Responses evoked by near-threshold intensities (blue) always had long latencies (>75 ms), but small increments in intensity (orange and green) evoked responses whose latencies were bimodally distributed. Further increments in intensity (red and purple) caused a near-complete switch to short-latency (<75 ms) responses. The proportion of slow and fast responses varied significantly with photostimulus intensity (χ^2^ = 105.0, *p* < 0.0001, excluding purple data points). Browne et al. (2017) reported a similar bimodal distribution of latencies but did not relate this to photostimulus intensity; instead, fast or slow responses occurred randomly in their experiments, perhaps because their ultra-short pulses (3 ms) activated neurons more variably, despite their high intensity (47 mW/mm^2^, which is >100× stronger than thresholds we measured using 100 ms-long pulses). Our data show that response latencies within each group decreased with increasing photostimulus intensity, as described by linear regressions on log-transformed data; the exponential drops in latency are clearest on the untransformed data (inset). One putative explanation, consistent with Browne et al. (2017) and with the double alarm system proposed by Plaghki et al. (2010) based on different rates of heating, is that slow and fast responses are mediated by C-and A-fibers, respectively. Building from that, our data suggest that C-fibers are recruited first (i.e. by weaker photostimuli) and that slow responses speed up as more C-fibers get recruited, but a discontinuous “switch” to fast responses occurs once A-fibers get recruited, and fast responses speed up as more A-fibers are recruited. Testing this requires further investigation, but resolving the stimulus-response relationship sufficiently to even pose such questions is notable.

To date, all reports of withdrawal from transcutaneous optogenetic stimulation used a pulse of blue light or a train of pulses (Abdo et al., 2019; Arcourt et al., 2017; Barik et al., 2018; Beaudry et al., 2017; Browne et al., 2017; Chamessian et al., 2019; Daou et al., 2013; Dhandapani et al., 2018; Iyer et al., 2014; Schorscher-Petcu et al., 2021; Sharif et al., 2020; Tashima et al., 2018; Warwick et al., 2021), with one exception, which used sustained light to activate keratinocytes (Baumbauer et al., 2015) (see also Discussion). Pulsed stimuli evoke precisely timed spikes (Browne et al., 2017), leading to spikes that are synchronized across co-activated neurons (Ratté et al., 2013), which may not accurately reflect the spiking patterns evoked by many types of somatosensory stimuli. That said, artificial stimuli need not mimic natural stimuli to be informative; indeed, deliberately evoking spiking patterns not possible with natural stimuli offers new opportunities to probe somatosensory coding (see Introduction). In that respect, optogenetic testing should not be limited to pulsed photostimuli. Capitalizing on the stability of our stimulator (see Fig. 2A), we therefore tested slowly ramped photostimuli for comparison with 100 ms-long pulses in two transgenic mouse lines (**Fig. 4B**): Advillin-ChR2 mice express ChR2 in all somatosensory afferents (Zhou et al., 2010) whereas Nav1.8-ChR2 mice express ChR2 selectively in nociceptors (Agarwal et al., 2004; Nassar et al., 2005). Both genotypes responded to pulses with bimodally distributed latencies, consistent with data in Figure 4A, but responses to ramps differed dramatically between genotypes, with all Nav1.8-ChR2 mice responding with a relatively short latency (1.1 ± 0.1 s; mean ± SD) whereas Advillin-ChR2 mice responded with much longer latencies on some trials. The difference was mostly due to intra-mouse variability rather than inter-mouse variability (i.e. individual Advillin-ChR2 mice responded with a mix of slow and fast responses rather than some mice being consistently slow and others being consistently fast). The basis for the genotypic difference requires further investigation though we hypothesize that co-activation of low-threshold mechanoreceptors in Advillin-ChR2 mice (but not in Nav1.8-ChR2 mice) engages a gate control mechanism that tempers the effects of nociceptive input, consistent with Arcourt et al. (2017), who showed that activating Aδ-HTMRs in isolation evoked more guarding, jumping and vocalization than co-activating Aδ-HTMRs and LTMRs. By testing different photostimulus waveforms, one can start to delineate the underlying interactions.

These results also raise questions about what stimulus parameter is most relevant. During ramps, many responses were initiated at intensities greater than the 5.74 mW used for the pulse. Channelrhodopsin-2 partially inactivates during sustained photostimulation but, despite this, the total charge passing through it by the time a ramp-evoked response occurs is dramatically higher than during a short pulse. Whether this is because abrupt-onset stimuli activate more neurons or activate them with a pattern that more readily triggers withdrawal requires further investigation, but clearly the photostimulus waveform is important.

### Other stimulation modalities and response measures

Radiant heat has long history of being used to quantify pain, from the Hardy-Wolff-Goodell dolorimeter for testing in humans (Hardy et al., 1940) to the Hargreaves test for rodents (Hargreaves et al., 1988). The latter remains popular in preclinical testing, second only to the von Frey test for mechanical sensitivity (Sadler et al., 2021). By incorporating an IR laser into the light path used for optogenetic stimulation, sensitivity to radiant heat can be tested using the same aiming and response measurement methods described above. To test our device, laser intensity was adjusted in pilot experiments to evoke withdrawal after ∼8 sec (like in a standard Hargreaves test) and stimuli were subsequently applied with a 20 s cut-off to avoid injury. Withdrawal latency was significantly reduced after injecting 0.5% capsaicin into the hind paw (**Fig. 5A**), as expected. Unlike existing stimulators which operate in close proximity to the paw, our device affords an unobstructed view of the paw from below. Substage video revealed that thermal stimulation triggered significantly more licking (**Fig. 5B**) and guarding (**Fig. 5C**) after capsaicin, which suggests that the heat stimulus is indeed more painful. Behaviors were manually scored here but this can be automated (**Fig. 5D** and **Video 6**). Automated classification in real-time would enable closed-loop control so that stimuli are applied contingent on a certain posture, e.g. when mice are standing on all four paws. Certain postures (e.g. guarding and rearing) can influence withdrawal latencies (Browne et al., 2017; Kauppila et al., 1998) and might therefore confound latency measurements.

**Figure 5.**
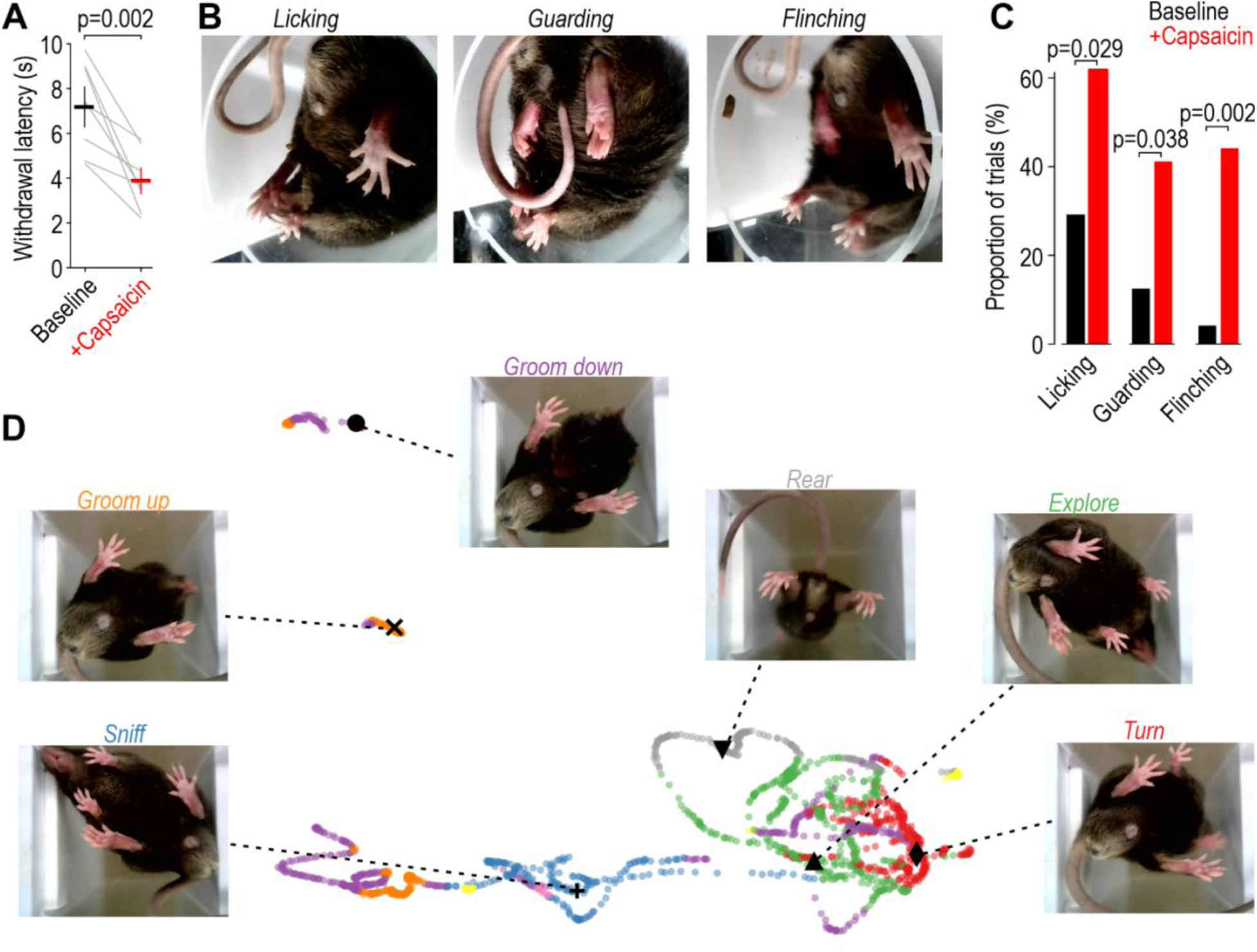
Multi-modal testing. (**A**) Radiant heat test. Latency dropped from 7.18 ± 0.74 s (mean ± SEM) at baseline to 3.88 ± 0.49 s after injecting 0.5% capsaicin into the left hind paw (*T*_7_ = 4.64, *p* = 0.002, paired *t*-test). (B) Associated behaviors. Sample frames illustrating licking, guarding, and flinching shortly after thermal stimulation (**Video 5**). **(C)** Proportion of trials in which these behaviors were exhibited in post-stimulus period increased significantly after capsaicin (*p* values on graph report χ^2^ tests). **(D)** Automated, non-supervised classification of ongoing behavior. Colored dots show latent space embeddings of behavioral state determined at regular intervals from **Video 6**. Screenshots corresponding to labeled points illustrate specific behaviors.

### Fully automated testing

Next, we mounted the stimulator on linear actuators (**Fig. 6A**) so aiming could be controlled remotely by joystick (i.e. without the tester operating in close proximity to the mice) or automatically using AI. For the latter, a neural network was trained using DeepLabCut to recognize the paws and other points on the mouse viewed from below. The computer is then fed the substage video stream and DeepLabCut-Live (Kane et al., 2020) uses the trained network to position the photostimulator by minimizing *x*- and *y*-error signals so that the target paw is positioned in the crosshairs for stimulation (**Fig. 6B**). Stimulation is initiated once the paw has remained stationary for a minimum period. Using a third, longer actuator, the device is shifted to the neighboring mouse after completing a trial. By interleaving trials, other mice are tested during the inter-stimulus interval required for each mouse. The order of testing can easily be randomized, which is difficult for an experimenter to keep track of, but trivial for a computer. Standardized high-throughput testing without the potential errors, systematic differences (Chesler et al., 2002), and animal stress (Sorge et al., 2014) associated with human testers is thus realized.

**Figure 6.**
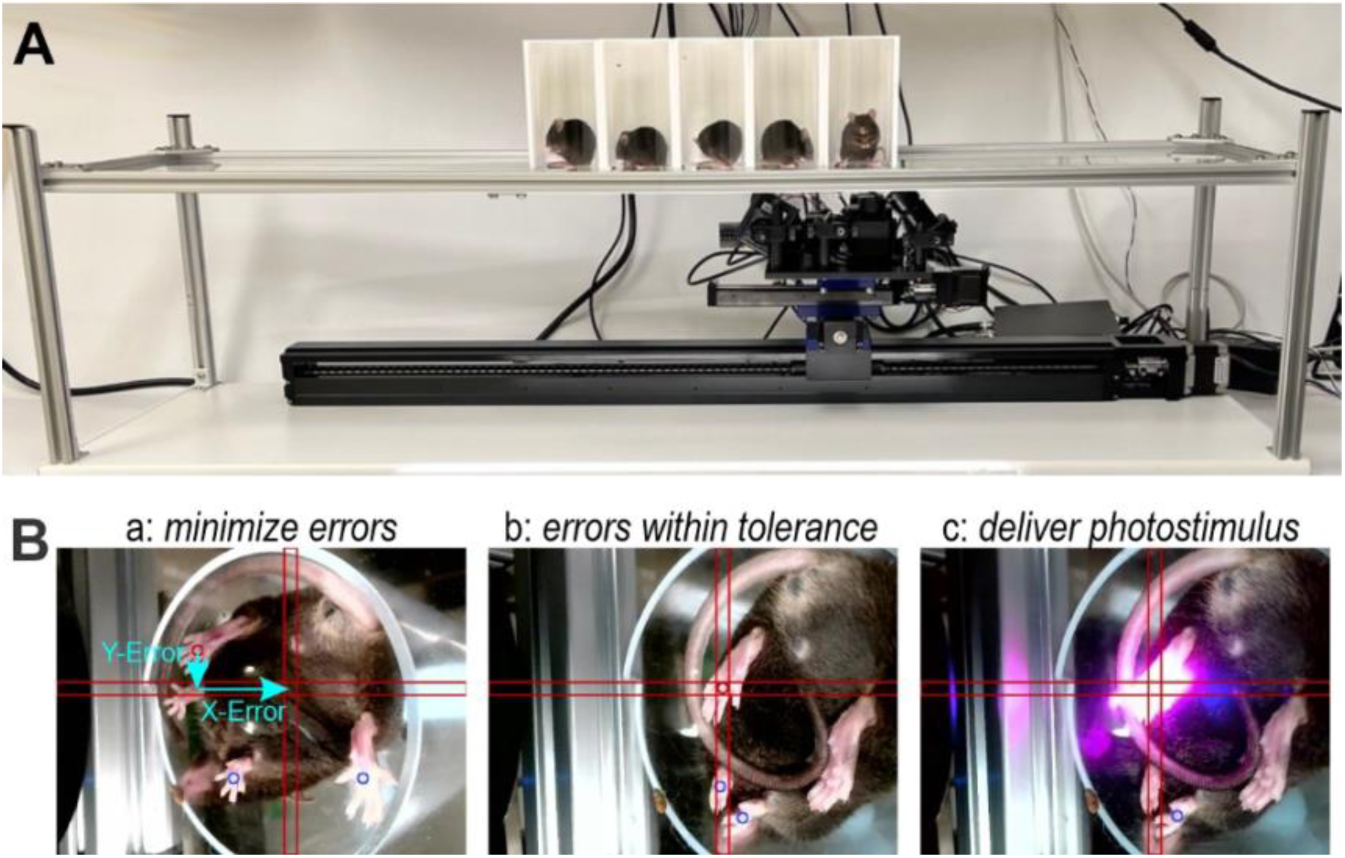
Fully automated testing. (**A**) Motorized stimulator. Unlike the manually aimed version (see Fig. 1), this version is mounted on linear actuators that allow for computer-controlled translation of the stimulator in the *x* and *y* axes. Aiming is controlled by joystick or automatically by AI. **(B)** Automated aiming. A neural network identifies the paws, snout and tail-base from the substage video. Deviation of the target paw from the crosshairs is measured (a) and then minimized by translating the stimulator (b). Once the paw has remained stable inside the crosshairs, photostimulation is initiated (c) and withdrawal latency is measured and recorded. The stimulator then moves to the adjacent mouse. See **Video 7** for example of fully automated testing with radiant heat.

## DISCUSSION

We developed a device able to reproducibly deliver photostimuli of different wavelengths, intensities, and kinetics (waveforms). This is a significant improvement over the handheld fiber optics previously used for transcutaneous optogenetic stimulation. We also incorporated a low-cost, high-speed photometer to detect paw withdrawal and measure withdrawal latency with millisecond precision based on changes in the reflectance of red light. The accuracy of this approach was validated by comparison with high-speed video. Closed-loop control of stimulation based on automated detection of reflectance changes is made possible by real-time data processing. Building on computer-controlled stimulation and response measurement, we automated the aiming process by training a neural network to recognize the paw and aim the stimulator at it using motorized actuators. Whereas aiming by joystick prevents the tester from working in close proximity to the mice, which stresses them (Sorge et al., 2014), automating the process removes the human element altogether, with significant benefits in terms of standardization, objectivity, and throughput.

Capitalizing on identification of gene expression patterns to distinguish subtypes of somatosensory afferents (Usoskin et al., 2015), optogenetics affords an unprecedented opportunity to study somatosensory coding by activating or inhibiting specific afferent subtypes. Testing the behavioral response to synthetic activation patterns allows one to explore causal relationships, complementing efforts to characterize activation patterns evoked by natural stimuli (Prescott et al., 2014). This does not require that optogenetic stimuli mimic natural, somatosensory stimuli. Optogenetic stimuli are patently unnatural – and the evoked sensations probably feel unnatural, like paresthesias evoked by electrical stimulation – but their ability to evoke behavior allows one to start inferring how they are perceived, and how those sensations relate to neural activation patterns. Doing this requires tight control of the stimulus and precise measurement of the response.

Technical advances have been made in delivering photostimuli to the CNS or peripheral nerves for optogenetic manipulations (Mickle et al., 2019). This technology is invaluable, but stimulating the nerve does not reproduce somatotopically organized stimulation. Photostimulating receptive fields in the skin is preferable in that regard, and is of course also less invasive. However, transcutaneous stimuli are difficult to apply reproducibly to behaving animals. Sharif et al. (2020) solved this by mounting the fiber optic to the head in order to stimulate the cheek, but a comparable solution is less feasible for stimulating the paw. These technical challenges explain why past studies focused on whether mice responded to optogenetic stimulation, without carefully varying stimulus parameters or measuring subtler aspects of the response. Past studies have varied the number, rate, or intensity of pulses, but in the suprathreshold regime, with consequences for the amount of licking, jumping, or vocalization. To our knowledge, only one previous study (Iyer et al., 2016) titrated the intensity of transcutaneous photostimuli to determine threshold (see Fig. 2D), and another (Schorscher-Petcu et al., 2021) titrated pulse duration and spot size. Moreover, scoring responses by eye, though still the norm for many tests, must be replaced with more objective, standardized metrics. Schorscher-Petcu et al. (2021) recently described a device that uses galvanometric mirrors to direct photostimuli and high-speed substage video to measure withdrawal (Schorscher-Petcu et al., 2021). Their device is very elegant but reliance on high-speed video to detect withdrawals likely precludes closed-loop control, and nor is their device fully automated or high-throughput. Our device delivers reproducible photostimuli and automatically measures withdrawals using the red-reflectance signal complemented by regular-speed substage video, which is also used for automated aiming.

Notably, non-painful stimuli may trigger withdrawal, which is to say that the threshold stimulus (or the probability of withdrawal) may not reflect painfulness (Jones et al., 2020). In that respect, testing with stronger stimuli is also informative. The study by Browne et al. (2017) stands out for its use of high-speed video to thoroughly quantify responses to optogenetic stimulation. Like us (see Fig. 4A), they observed a bimodal distribution of withdrawal latencies; however, they observed this despite using high-intensity pulses, most likely because their pulses were extremely brief (3 ms) and might, therefore, have activated afferents probabilistically. Browne et al. (2017) also noted that the withdrawal response was not limited to the stimulated limb, and instead involved a more widespread motor response. Though not quantified here, a widespread response was evident in the substage video, and sometimes including vocalizing, facial grimacing, jumping, and orienting to the stimulus followed by licking, guarding or flinching of the stimulated paw (see Fig. 5B,C). A complete analysis of each trial ought to consider not only the reflexive component (i.e. did withdrawal occur and how quickly), but also whether signs of discomfort were exhibited afterwards and for how long. Those signs are obvious when applying very strong stimuli, but become harder to discern with near-threshold stimulation, which makes objective quantification all the more important. Regular-speed video is sufficient to capture all but the fast reflex (which can be measured by other means; see Fig. 3) and the bottom-up view is well suited for AI-based quantification of ongoing behaviors (Hsu & Yttri, 2021; Luxem et al., 2022). We recommend that video be recorded for all trials if only to allow retrospective analysis of those data in the future. There has been an explosion of AI-based methods for extracting key points on animals (Graving et al., 2019; Mathis et al., 2018; Pereira et al., 2022) and algorithms for extracting higher-level animal behaviors from key point (Hsu & Yttri, 2021; Luxem et al., 2022; Weinreb et al., 2023) or raw pixel (Bohnslav et al., 2021) data, and rapid advances are likely to continue. By capturing video before and after all stimuli, users can incorporate their own AI-based methods or manually score behaviors for consideration above and beyond reflex measurements.

Withdrawal responses are known to be sensitive to the rodent’s posture and ongoing behavior at the time of stimulation (Blivis et al., 2017; Browne et al., 2017; Kauppila et al., 1998). On the one hand, such differences may confound latency measurements but, on the other hand, they may provide important information; either way, they should be accounted for. Waiting for each mouse to adopt a specific posture and behavioral state is onerous for human testers but is something that a neural network can be trained to do, with automated stimulation being made contingent on the mouse exhibiting a certain state. But before that, to better understand the relationship between the mouse’s pre-stimulus state and its subsequent stimulus-evoked withdrawal (and post-stimulus state), the state at the onset of stimulation could be classified from video (like in Fig. 5D) and correlated with the evoked response on that trial. Other comparisons would also be informative, like correlating the pre-and post-stimulus states and treatment status. In short, more data can be acquired and more thoroughly analyzed than is typically done in current protocols. This need not entail expensive equipment or reduced throughput.

Nearly all past studies involving transcutaneous optogenetic stimulation used single pulses or pulse trains. In the one exception, Baumbauer et al. (2015) activated ChR2-expressing keratinocytes with sustained light. Single pulses and pulse trains are just some of the many possible waveforms, especially since LEDs can be so easily controlled. Notably, just like pulsed electrical stimuli lost favor in pain testing because of the unnaturally synchronized neural activation they evoke (Le Bars et al., 2001), pulsed optogenetic stimuli warrant similar scrutiny and should not be the only waveform tested. Indeed different rates of radiant heating differentially engage C and A fibers, thus enabling the role of different afferents to be studies (Plaghki et al., 2010). By testing photostimulus ramps, we uncovered genotypic differences that were not evident with pulses (see Fig. 4B). In addition to transcutaneous optogenetic pulses to single paws, some studies (Dao et al. 2013; Iyer et al. 2014) have tested if ChR2-expressing mice avoid blue-lit floors, which they do. In these cases, the floor light was continuous, unlike the pulses applied by fiber optic; it is, therefore, notable that mice avoided the blue floor but did not respond to it with reflexive withdrawal, paw licking, or other outward signs of pain, like they did to pulses. In another case where the floor light was pulsed (Barik et al., 2018), reflexive withdrawal was observed. These results highlight the underappreciated importance of the stimulus waveform.

To summarize, we describe a new device capable of reproducible stimulation and precise yet efficient response measurement. Aiming, stimulation, and measurement can be fully automated, which improves standardization and increases throughput, amongst other benefits. A video record of the animal before, during and after stimulation allows one to extend analysis beyond traditional response metrics (i.e. threshold and latency) to consider if/how evoked and spontaneous pain behaviors are correlated.

## MATERIALS AND METHODS

### Animals

All procedures were approved by the Animal Care Committee at The Hospital for Sick Children (protocol #53451) and were conducted in accordance with guidelines from the Canadian Council on Animal Care. To express ChR2 selectively in different types of primary somatosensory afferents, we used Ai32(RCL-ChR2(H134R)/EYFP) mice (JAX:024109), which express the H134R variant of ChR2 in cells expressing Cre recombinase. These were crossed with advillin^Cre^ mice (kindly provided by Fan Wang) to express ChR2 in all sensory afferents, TrpV1^Cre^ mice (JAX:017769) to express ChR2 in TrpV1-lineage neurons, or Na_V_1.8^Cre^ mice (kindly provided by Rohini Kuner) to express ChR2 in nociceptors. Mice were acclimated to their testing chambers for 1 hr on the day before the first day of testing, and each day for 1 hr prior to the start of testing.

### Photostimulator

The stimulator is summarized in **Supplementary Figure 1**; A complete list of components (as numbered in Supplementary Fig. 1, with part # and supplier information) is included as **Supplementary Table 1**. Briefly, collimated light from a red (625 nm) LED and blue (455 nm) LED is combined using a 550 nm cut-on dichroic mirror. Blue light is attenuated with a neutral density filter. This beam is combined with IR light from a 980 nm solid laser using a 900 nm cut-on dichroic mirror. The IR beam is expanded to fill the back of the focusing lens. The common light path is reflected upward with a mirror and focussed to a spot 5 mm in diameter on the platform above. The surface area of the spot is ∼20 mm^2^; photostimulus intensity values should be divided by this number to convert to light density (irradiance). Red light reflected off the mouse paw is collected by a photodetector through a 630 nm notch filter. All light sources are controlled by computer via appropriate drivers and a 1401 DAQ (Cambridge Electronic Design) using Spike2 (Cambridge Electronic Design) or custom software written in Python. The photodetector samples at 1 kHz with the same DAQ, thus synchronizing stimulation and withdrawal measurement. A borescope provides video of the mouse from below (substage). Video is used for aiming with the help of visual feedback using the red light, which is turned on prior to photostimulation with blue or IR light. A near-IR light source is useful to improve lighting during high-speed video. In the manual version of the device, the device is slid by hand; leveling screws at the four corners of the breadboard have a plastic cap for smooth sliding. In the motorized version, the breadboard it attached to linear actuators (TBI Motion) via 3-D printed connectors. Motors are controlled via custom software. The user aims by keyboard or joystick, or fully automated aiming is left to a neural network trained to recognize the mouse paws (see Results). All measurements of light delivery were taken using a PM100D optical power meter with a s170C photodiode (Thorlabs).

### Platform and enclosures

The platform and animal enclosures were custom made. The platform is 3 mm-thick clear Plexiglass mounted on 20×20 mm aluminum rails, adjusted to the desired height above the stimulator. Various enclosure designed were tested. In the final design, clear Plexiglass tubes (outer diameter = 65 mm, thickness = 2 mm) cut in 12.5 cm lengths were used in conjunction with opaque white 3-D printed cubicle. The same tube used to transfer a mouse from its home cage is placed on the platform vertically and slid into a cubicle for testing (see Fig. 1B). A notch cut into the base of each tube allows the experimenter to deliver a food reward, to poke the mouse (to wake or orient it), or to clean feces or urine from the platform if required. Each cubicle is 3-D printed and contains internal magnets that allow cubicles to be easily combined. Keeping the mice at fixed distances from each other is important for automated testing, where the stimulator is automatically translated a fixed distance when testing consecutive mice. To view the mouse in profile during high-speed video, we used a narrow rectangular chamber with clear walls on the front and left side (with a notch under the latter) and opaque walls at the rear and right side. In some cases, a mirror was placed at a 45° angle near the left wall to simultaneously capture a front view of the mouse.

### Automated withdrawal detection and latency measurement

Paw withdrawal is detected and its latency measured from the red reflectance signal using custom code written in Python. Red light is initiated prior to photostimulation with blue or IR light. Baseline reflectance is measured over the 0.5 s epoch preceding photostimulus onset. A running average across a 27 ms-wide window was used to remove noise. Withdrawal latency was defined as time elapsed from photostimulus onset until the reflectance signal dropped below a threshold defined as 2 mV below baseline; the signal needed to remain below threshold for >20 ms to qualify as a response, but latency was calculated based on the start of that period. Latencies thus extracted from the reflectance signal were compared to latency values extracted from high-speed video of the same withdrawal. In the latter case, paw height was extracted from video (see below) using DeepLabCut; latency was taken as the time taken for paw height to rise 6 pixels above baseline, defined as the mean height over the 0.5 s epoch preceding photostimulus onset,

### High-speed video

High-speed video was collected with a Chronos 1.4 camera (Krontech) using a Computar 12.5-75 mm f/1.2 lens sampling at 1000 fps. To synchronize video with stimulation, the camera was triggered with digital pulses sent from the DAQ. Videos were compressed using H.264. Video was analyzed using DeepLabCut (Mathis et al., 2018) to label the hind paw in sample frames and train a deep neural network to recognize the paw. This returned paw trajectories which were analyzed using custom code written in Python.

### Substage Behavior Extraction

Substage video was saved and compressed using H.264. DeepLabCut was used to identify the nose, left fore paw, right fore paw, left hind paw, right hind paw, and tail base. These key points were then fed into the VAME framework to extract complex behaviors (Luxem et al., 2022).

### Capsaicin and heat hypersensitivity

A 0.5% w/v solution of capsaicin in mineral oil was prepared. Mice were lightly anesthetized using isofluorane and 5 µL of capsaicin was injected into the left hind paw. The mice were allowed to recover for 15 min from anesthesia before thermal sensitivity was assessed.

## Acknowledgements

This study was funded by a Foundation Grant from the Canadian Institutes of Health Research (FDN167276) and a Proof-of-Principle grant from The Hospital for Sick Children to SAP. We thank Jason Jeong for expert technical assistance, Erica Austriaco for contributions to the software, and volunteer testers whose participation allowed us to compare light delivery methods.

## SUPPLEMENTARY MATERIAL

Supplementary Table 1. Complete list of components, including sources and part numbers.

Video 1. Red light during aiming and paw withdrawal.

Video 2a. Measurement of light delivery using light meter – Handheld fiber optic.

Video 2b. Measurement of light delivery using light meter – Photostimulator.

Video 3. High-speed video of mouse in profile to measure paw height.

Video 4. Responses to photostimulus pulse and ramp

Video 5a. Responses to radiant heat before intraplantar capsaicin.

Video 5b. Responses to radiant heat after intraplantar capsaicin.

Video 6. Ongoing behavior and its automated classification.

Video 7. Example of fully automated testing with radiant heat and graphical user interface (GUI) of associated software.

**Supplementary Figure 1.**
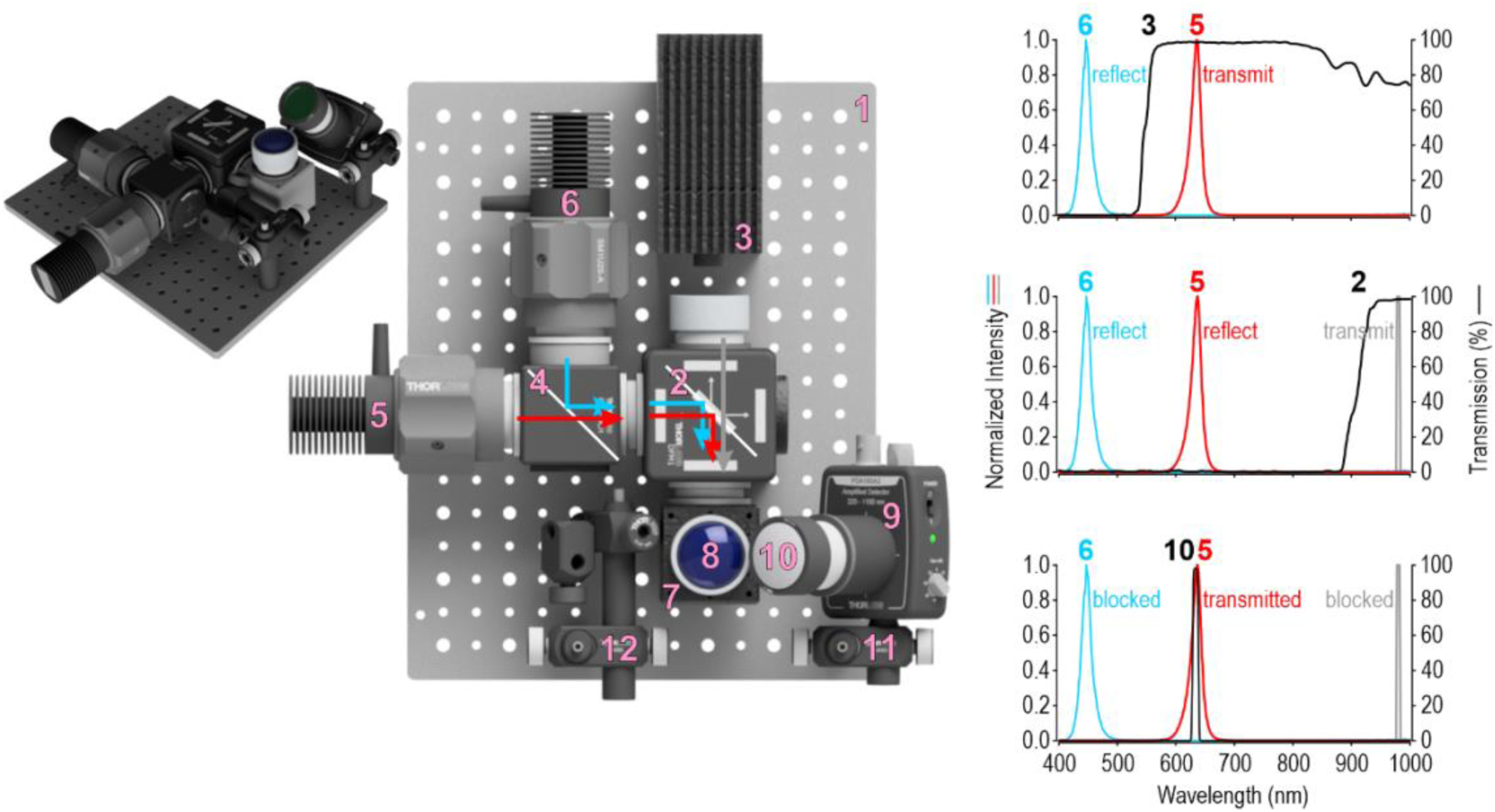
Stimulator components. Key components of the stimulator are numbered *1-12*, with details provided in the corresponding entries in Table S1. Emission spectra for the light sources (*3, 5, 6*) are shown relative to the transmission properties of the dichroic mirrors (*2, 4*) and notch filter (*10*) to appreciate how lights sources are combined, and what light is measured by the photodetector (*9*). All curves except for IR laser are based on data provided by ThorLabs. IR laser is reported to produce 980±5 nm light.

**Supplementary Figure 2.**
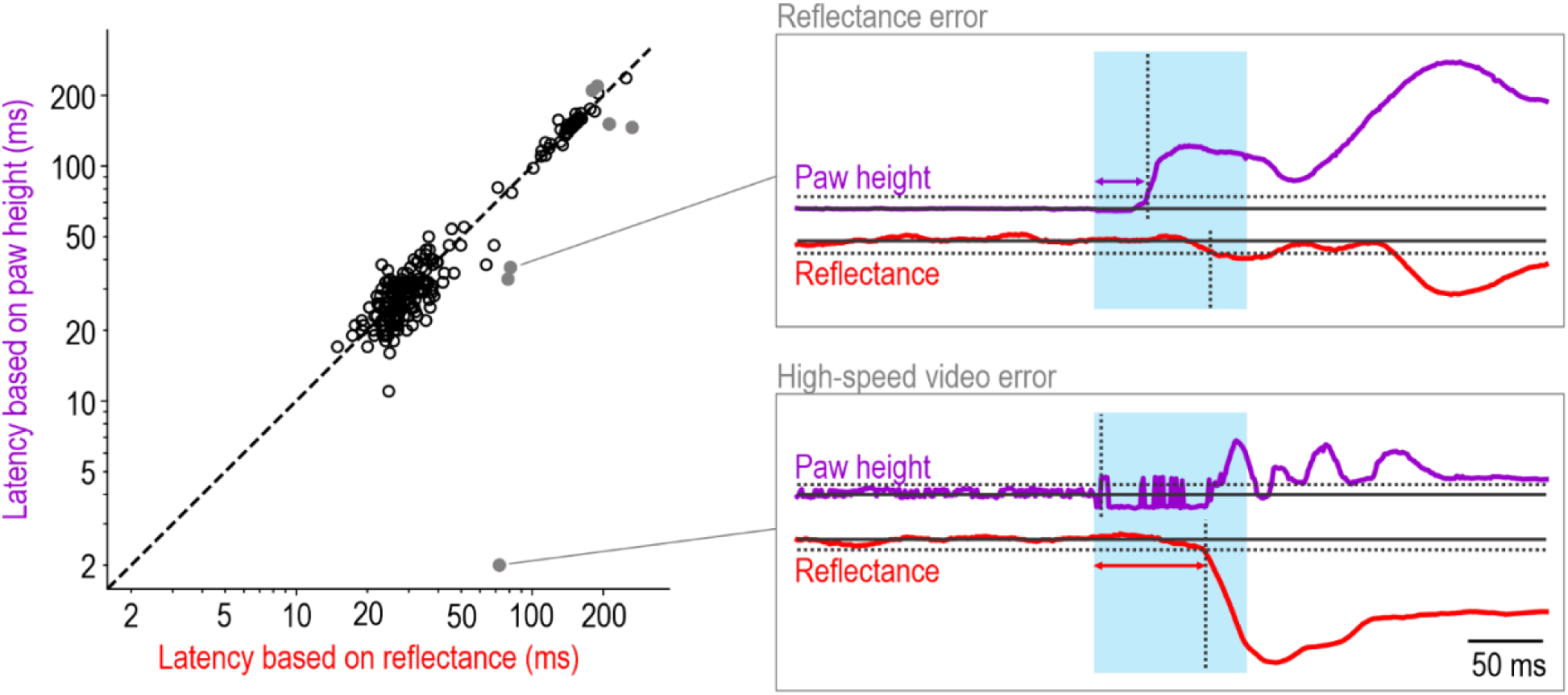
Errors in withdrawal latency measurement. Data are plotted like in Figure 2B but errors trials are now included as grey dots. Of the seven error trials identified through visual inspection of all trials, automated determination of paw position from high-speed video was corrupted by the blue light during photostimulation in 3 trials (top) and the reflectance signal did not immediately change upon paw withdrawal in the other 4 trials (bottom). The false negative rate for the reflectance signal is thus <2% and we did not identify any false positives (i.e. changes in reflectance in the absence of paw movement).

## REFERENCES

Abboud, C., Duveau, A., Bouali-Benazzouz, R., Massé, K., Mattar, J., Brochoire, L., Fossat, P., Boué-Grabot, E., Hleihel, W., & Landry, M. (2021). Animal models of pain: Diversity and benefits. Journal of Neuroscience Methods, 348, 108997. https://doi.org/10.1016/J.JNEUMETH.2020.108997

Abdo, H., Calvo-Enrique, L., Lopez, J. M., Song, J., Zhang, M. D., Usoskin, D., Manira, A. El, Adameyko, I., Hjerling-Leffler, J., & Ernfors, P. (2019). Specialized cutaneous schwann cells initiate pain sensation. Science, 365(6454), 695–699. https://doi.org/10.1126/SCIENCE.AAX6452

Abdus-Saboor, I., Fried, N. T., Lay, M., Burdge, J., Swanson, K., Fischer, R., Jones, J., Dong, P., Cai, W., Guo, X., Tao, Y. X., Bethea, J., Ma, M., Dong, X., Ding, L., & Luo, W. (2019). Development of a Mouse Pain Scale Using Sub-second Behavioral Mapping and Statistical Modeling. Cell Reports, 28(6), 1623–1634.e4. https://doi.org/10.1016/j.celrep.2019.07.017

Agarwal, N., Offermanns, S., & Kuner, R. (2004). Conditional gene deletion in primary nociceptive neurons of trigeminal ganglia and dorsal root ganglia. Genesis, 38(3), 122–129. https://doi.org/10.1002/GENE.20010

Arcourt, A., Gorham, L., Dhandapani, R., Prato, V., Taberner, F. J., Wende, H., Gangadharan, V., Birchmeier, C., Heppenstall, P. A., & Lechner, S. G. (2017). Touch Receptor-Derived Sensory Information Alleviates Acute Pain Signaling and Fine-Tunes Nociceptive Reflex Coordination. Neuron, 93(1), 179–193. https://doi.org/10.1016/j.neuron.2016.11.027

Arnold, L. M., Bennett, R. M., Crofford, L. J., Dean, L. E., Clauw, D. J., Goldenberg, D. L., Fitzcharles, M. A., Paiva, E. S., Staud, R., Sarzi-Puttini, P., Buskila, D., & Macfarlane, G. J. (2019). AAPT Diagnostic Criteria for Fibromyalgia. The Journal of Pain, 20(6), 611–628. https://doi.org/10.1016/J.JPAIN.2018.10.008

Backonja, M. M., & Stacey, B. (2004). Neuropathic pain symptoms relative to overall pain rating. The Journal of Pain, 5(9), 491–497. https://doi.org/10.1016/J.JPAIN.2004.09.001

Barik, A., Thompson, J. H., Seltzer, M., Ghitani, N., & Chesler, A. T. (2018). A Brainstem-Spinal Circuit Controlling Nocifensive Behavior. Neuron, 100(6), 1491–1503.e3. https://doi.org/10.1016/J.NEURON.2018.10.037

Baron, R., Dickenson, A. H., Calvo, M., Dib-Hajj, S. D., & Bennett, D. L. (2022). Maximizing treatment efficacy through patient stratification in neuropathic pain trials. Nature Reviews Neurology, 19, 53–64. https://doi.org/10.1038/S41582-022-00741-7

Barrot, M. (2012). Tests and models of nociception and pain in rodents. Neuroscience, 211, 39–50. https://doi.org/10.1016/J.NEUROSCIENCE.2011.12.041

Baumbauer, K. M., Deberry, J. J., Adelman, P. C., Miller, R. H., Hachisuka, J., Lee, K. H., Ross, S. E., Koerber, H. R., Davis, B. M., & Albers, K. M. (2015). Keratinocytes can modulate and directly initiate nociceptive responses. ELife, 4(September 2015). https://doi.org/10.7554/ELIFE.09674

Beaudry, H., Daou, I., Ase, A. R., Ribeiro-Da-Silva, A., & Séguela, P. (2017). Distinct behavioral responses evoked by selective optogenetic stimulation of the major TRPV1+ and MrgD+ subsets of C-fibers. Pain, 158(12), 2329–2339. https://doi.org/10.1097/J.PAIN.0000000000001016

Blivis, D., Haspel, G., Mannes, P. Z., O’Donovan, M. J., & Iadarola, M. J. (2017). Identification of a novel spinal nociceptive-motor gate control for Aδ pain stimuli in rats. ELife, 6. https://doi.org/10.7554/ELIFE.23584

Bohnslav, J. P., Wimalasena, N. K., Clausing, K. J., Dai, Y. Y., Yarmolinsky, D. A., Cruz, T., Kashlan, A. D., Chiappe, M. E., Orefice, L. L., Woolf, C. J., & Harvey, C. D. (2021). DeepEthogram, a machine learning pipeline for supervised behavior classification from raw pixels. ELife, 10. https://doi.org/10.7554/ELIFE.63377

Browne, L. E., Latremoliere, A., Lehnert, B. P., Grantham, A., Ward, C., Alexandre, C., Costigan, M., Michoud, F., Roberson, D. P., Ginty, D. D., & Woolf, C. J. (2017). Time-Resolved Fast Mammalian Behavior Reveals the Complexity of Protective Pain Responses. Cell Reports, 20(1), 89–98. https://doi.org/10.1016/j.celrep.2017.06.024

Burma, N. E., Leduc-Pessah, H., Fan, C. Y., & Trang, T. (2017). Animal models of chronic pain: Advances and challenges for clinical translation. Journal of Neuroscience Research, 95(6), 1242–1256. https://doi.org/10.1002/JNR.23768

Chamessian, A., Matsuda, M., Young, M., Wang, M., Zhang, Z. J., Liu, D., Tobin, B., Xu, Z. Z., Van de Ven, T., & Ji, R. R. (2019). Is Optogenetic Activation of Vglut1-Positive Aβ Low-Threshold Mechanoreceptors Sufficient to Induce Tactile Allodynia in Mice after Nerve Injury? The Journal of Neuroscience, 39(31), 6202–6215. https://doi.org/10.1523/JNEUROSCI.2064-18.2019

Chesler, E. J., Wilson, S. G., Lariviere, W. R., Rodriguez-Zas, S. L., & Mogil, J. S. (2002). Influences of laboratory environment on behavior. Nature Neuroscience, 5(11), 1101–1102. https://doi.org/10.1038/NN1102-1101

Copits, B. A., Pullen, M. Y., & Gereau, R. W. (2016). Spotlight on pain: Optogenetic approaches for interrogating somatosensory circuits. Pain, 157(11), 2424–2433. https://doi.org/10.1097/J.PAIN.0000000000000620

Daou, I., Tuttle, A. H., Longo, G., Wieskopf, J. S., Bonin, R. P., Ase, A. R., Wood, J. N., De Koninck, Y., Ribeiro-da-Silva, A., Mogil, J. S., & Séguéla, P. (2013). Remote Optogenetic Activation and Sensitization of Pain Pathways in Freely Moving Mice. The Journal of Neuroscience, 33(47), 18631. https://doi.org/10.1523/JNEUROSCI.2424-13.2013

De Farias Rocha, F. A., Gomes, B. D., De Lima Silveira, L. C., Martins, S. L., Aguiar, R. G., De Souza, J. M., & Ventura, D. F. (2016). Spectral Sensitivity Measured with Electroretinogram Using a Constant Response Method. PLOS ONE, 11(1), e0147318. https://doi.org/10.1371/JOURNAL.PONE.0147318

Deuis, J. R., Dvorakova, L. S., & Vetter, I. (2017). Methods used to evaluate pain behaviors in rodents. In Frontiers in Molecular Neuroscience (Vol. 10, p. 284). Frontiers Media S.A. https://doi.org/10.3389/fnmol.2017.00284

Dhandapani, R., Arokiaraj, C. M., Taberner, F. J., Pacifico, P., Raja, S., Nocchi, L., Portulano, C., Franciosa, F., Maffei, M., Hussain, A. F., De Castro Reis, F., Reymond, L., Perlas, E., Garcovich, S., Barth, S., Johnsson, K., Lechner, S. G., & Heppenstall, P. A. (2018). Control of mechanical pain hypersensitivity in mice through ligand-targeted photoablation of TrkB-positive sensory neurons. Nature Communications, 9(1640), 1–14. https://doi.org/10.1038/s41467-018-04049-3

Edwards, R. R., Dworkin, R. H., Turk, D. C., Angst, M. S., Dionne, R., Freeman, R., Hansson, P., Haroutounian, S., Arendt-Nielsen, L., Attal, N., Baron, R., Brell, J., Bujanover, S., Burke, L. B., Carr, D., Chappell, A. S., Cowan, P., Etropolski, M., Fillingim, R. B., … Yarnitsky, D. (2016). Patient phenotyping in clinical trials of chronic pain treatments: IMMPACT recommendations. Pain, 157(9), 1851–1871. https://doi.org/10.1097/J.PAIN.0000000000000602

Fried, N. T., Chamessian, A., Zylka, M. J., & Abdus-Saboor, I. (2020). Improving pain assessment in mice and rats with advanced videography and computational approaches. PAIN, 161(7).

Gouveia, K., & Hurst, J. L. (2013). Reducing Mouse Anxiety during Handling: Effect of Experience with Handling Tunnels. PLoS ONE, 8(6), 66401. https://doi.org/10.1371/JOURNAL.PONE.0066401

Gouveia, K., & Hurst, J. L. (2019). Improving the practicality of using non-aversive handling methods to reduce background stress and anxiety in laboratory mice. Scientific Reports 2019 9:1, *9*(1), 1–19. https://doi.org/10.1038/s41598-019-56860-7

Graving, J. M., Chae, D., Naik, H., Li, L., Koger, B., Costelloe, B. R., & Couzin, I. D. (2019). Deepposekit, a software toolkit for fast and robust animal pose estimation using deep learning. ELife, 8. https://doi.org/10.7554/ELIFE.47994

Gregory, N. S., Harris, A. L., Robinson, C. R., Dougherty, P. M., Fuchs, P. N., & Sluka, K. A. (2013). An overview of animal models of pain: disease models and outcome measures. The Journal of Pain, 14(11), 1255–1269. https://doi.org/10.1016/J.JPAIN.2013.06.008

Hardy, J. D., Wolff, H. G., & Goodell, H. (1940). STUDIES ON PAIN. A NEW METHOD FOR MEASURING PAIN THRESHOLD: OBSERVATIONS ON SPATIAL SUMMATION OF PAIN. The Journal of Clinical Investigation, 19(4), 649–657. https://doi.org/10.1172/JCI101168

Hargreaves, K., Dubner, R., Brown, F., Flores, C., & Joris, J. (1988). A new and sensitive method for measuring thermal nociception in cutaneous hyperalgesia. Pain, 32(1), 77–88. https://doi.org/10.1016/0304-3959(88)90026-7

Hsu, A. I., & Yttri, E. A. (2021). B-SOiD, an open-source unsupervised algorithm for identification and fast prediction of behaviors. Nature Communications 2021 12:1, *12*(1), 1–13. https://doi.org/10.1038/S41467-021-25420-X

Hurst, J. L., & West, R. S. (2010). Taming anxiety in laboratory mice. Nature Methods, 7(10), 825–826. https://doi.org/10.1038/nmeth.1500

Iyer, S. M., Montgomery, K. L., Towne, C., Lee, S. Y., Ramakrishnan, C., Deisseroth, K., & Delp, S. L. (2014). Virally mediated optogenetic excitation and inhibition of pain in freely moving nontransgenic mice. Nature Biotechnology 2014 32:3, *32*(3), 274–278. https://doi.org/10.1038/NBT.2834

Iyer, S. M., Vesuna, S., Ramakrishnan, C., Huynh, K., Young, S., Berndt, A., Lee, S. Y., Gorini, C. J., Deisseroth, K., & Delp, S. L. (2016). Optogenetic and chemogenetic strategies for sustained inhibition of pain. Scientific Reports 2016 6:1, *6*(1), 1–10. https://doi.org/10.1038/SREP30570

Jaggi, A. S., Jain, V., & Singh, N. (2011). Animal models of neuropathic pain. Fundamental & Clinical Pharmacology, 25(1), 1–28. https://doi.org/10.1111/J.1472-8206.2009.00801.X

Jones, J. M., Foster, W., Twomey, C. R., Burdge, J., Ahmed, O. M., Pereira, T. D., Wojick, J. A., Corder, G., Plotkin, J. B., & Abdus-Saboor, I. (2020). A machine-vision approach for automated pain measurement at millisecond timescales. ELife, 9. https://doi.org/10.7554/eLife.57258

Kane, G. A., Lopes, G., Saunders, J. L., Mathis, A., & Mathis, M. W. (2020). Real-time, low-latency closed-loop feedback using markerless posture tracking. ELife, 9, 1–29. https://doi.org/10.7554/ELIFE.61909

Kauppila, T., Kontinen, V. K., & Pertovaara, A. (1998). Weight bearing of the limb as a confounding factor in assessment of mechanical allodynia in the rat. Pain, 74(1), 55–59. https://doi.org/10.1016/S0304-3959(97)00143-7

Koltzenburg, M., Torebjörk, H. E., & Wahren, L. K. (1994). Nociceptor modulated central sensitization causes mechanical hyperalgesia in acute chemogenic and chronic neuropathic pain. Brain, 117(3), 579–591. https://doi.org/10.1093/BRAIN/117.3.579

Le Bars, D., Gozariu, M., & Cadden, S. W. (2001). Animal Models of Nociception. Pharmacological Reviews, 53(4), 597.

Le Bars, D., Hansson, P. T., & Plaghki, L. (2009). Current animal test and models of pain. In *Pharmacology of Pain* (pp. 475–504).

Luxem, K., Mocellin, P., Fuhrmann, F., Kürsch, J., Miller, S. R., Palop, J. J., Remy, S., & Bauer, P. (2022). Identifying behavioral structure from deep variational embeddings of animal motion. Communications Biology 2022 5:1, *5*(1), 1–15. https://doi.org/10.1038/S42003-022-04080-7

Maier, C., Baron, R., Tölle, T. R., Binder, A., Birbaumer, N., Birklein, F., Gierthmühlen, J., Flor, H., Geber, C., Huge, V., Krumova, E. K., Landwehrmeyer, G. B., Magerl, W., Maihöfner, C., Richter, H., Rolke, R., Scherens, A., Schwarz, A., Sommer, C., … Treede, R. D. (2010). Quantitative sensory testing in the German Research Network on Neuropathic Pain (DFNS): Somatosensory abnormalities in 1236 patients with different neuropathic pain syndromes. Pain, 150(3), 439–450. https://doi.org/10.1016/J.PAIN.2010.05.002

Mathis, A., Mamidanna, P., Cury, K. M., Abe, T., Murthy, V. N., Mathis, M. W., & Bethge, M. (2018). DeepLabCut: markerless pose estimation of user-defined body parts with deep learning. Nature Neuroscience 2018 21:9, *21*(9), 1281–1289. https://doi.org/10.1038/s41593-018-0209-y

Mickle, A. D., Won, S. M., Noh, K. N., Yoon, J., Meacham, K. W., Xue, Y., McIlvried, L. A., Copits, B. A., Samineni, V. K., Crawford, K. E., Kim, D. H., Srivastava, P., Kim, B. H., Min, S., Shiuan, Y., Yun, Y., Payne, M. A., Zhang, J., Jang, H., … Rogers, J. A. (2019). A wireless closed-loop system for optogenetic peripheral neuromodulation. Nature, 565(7739), 361–365. https://doi.org/10.1038/S41586-018-0823-6

Mogil, J. S. (2017). Laboratory environmental factors and pain behavior: the relevance of unknown unknowns to reproducibility and translation. Lab Animal, 46(4), 136–141. https://doi.org/10.1038/laban.1223

Mogil, J. S., & Crager, S. E. (2004). What should we be measuring in behavioral studies of chronic pain in animals? Pain, 112(1), 12–15. https://doi.org/10.1016/J.PAIN.2004.09.028

Mogil, J. S., Graham, A. C., Ritchie, J., Hughes, S. F., Austin, J. S., Schorscher-Petcu, A., Langford, D. J., & Bennett, G. J. (2010). Hypolocomotion, asymmetrically directed behaviors (licking, lifting, flinching, and shaking) and dynamic weight bearing (gait) changes are not measures of neuropathic pain in mice. Molecular Pain, 6. https://doi.org/10.1186/1744-8069-6-34

Nassar, M. A., Levato, A., Stirling, L. C., & Wood, J. N. (2005). Neuropathic pain develops normally in mice lacking both Nav 1.7 and Nav 1.8. Molecular Pain, 1. https://doi.org/10.1186/1744-8069-1-24

Negus, S. S. (2019). Core Outcome Measures in Preclinical Assessment of Candidate Analgesics. Pharmacological Reviews, 71(2), 225–266. https://doi.org/10.1124/PR.118.017210

Nikbakht, N., & Diamond, M. E. (2021). Conserved visual capacity of rats under red light. ELife, 10. https://doi.org/10.7554/ELIFE.66429

Niklaus, S., Albertini, S., Schnitzer, T. K., & Denk, N. (2020). Challenging a Myth and Misconception: Red-Light Vision in Rats. Animals, 10(3), 422. https://doi.org/10.3390/ANI10030422

Pereira, T. D., Tabris, N., Matsliah, A., Turner, D. M., Li, J., Ravindranath, S., Papadoyannis, E. S., Normand, E., Deutsch, D. S., Wang, Z. Y., McKenzie-Smith, G. C., Mitelut, C. C., Castro, M. D., D’Uva, J., Kislin, M., Sanes, D. H., Kocher, S. D., Wang, S. S. H., Falkner, A. L., … Murthy, M. (2022). SLEAP: A deep learning system for multi-animal pose tracking. Nature Methods 2022 19:4, *19*(4), 486–495. https://doi.org/10.1038/S41592-022-01426-1

Pitzer, C., Kuner, R., & Tappe-Theodor, A. (2016). Voluntary and evoked behavioral correlates in neuropathic pain states under different social housing conditions. Molecular Pain, 12. https://doi.org/10.1177/1744806916656635

Plaghki, L., Decruynaere, C., Van Dooren, P., & Le Bars, D. (2010). The Fine Tuning of Pain Thresholds: A Sophisticated Double Alarm System. PLOS ONE, 5(4), e10269. https://doi.org/10.1371/JOURNAL.PONE.0010269

Prescott, S. A., Ma, Q., & De Koninck, Y. (2014). Normal and abnormal coding of somatosensory stimuli causing pain. Nature Neuroscience, 17, 183–191. https://doi.org/10.1038/nn.3629

Ratté, S., Hong, S., DeSchutter, E., & Prescott, S. A. (2013). Impact of neuronal properties on network coding: Roles of spike initiation dynamics and robust synchrony transfer. In Neuron (Vol. 78, Issue 5, pp. 758–772). https://doi.org/10.1016/j.neuron.2013.05.030

Rowbotham, M. C., & Fields, H. L. (1996). The relationship of pain, allodynia and thermal sensation in post-herpetic neuralgia. Brain, 119(2), 347–354. https://doi.org/10.1093/BRAIN/119.2.347

Sadler, K. E., Mogil, J. S., & Stucky, C. L. (2021). Innovations and advances in modelling and measuring pain in animals. Nature Reviews Neuroscience, 23(2), 70–85. https://doi.org/10.1038/S41583-021-00536-7

Schorscher-Petcu, A., Takács, F., & Browne, L. E. (2021). Scanned optogenetic control of mammalian somatosensory input to map input-specific behavioral outputs. ELife, 10. https://doi.org/10.7554/ELIFE.62026

Sharif, B., Ase, A. R., Ribeiro-da-Silva, A., & Séguéla, P. (2020). Differential Coding of Itch and Pain by a Subpopulation of Primary Afferent Neurons. Neuron, 106(6), 940–951.e4. https://doi.org/10.1016/J.NEURON.2020.03.021

Sorge, R. E., Martin, L. J., Isbester, K. A., Sotocinal, S. G., Rosen, S., Tuttle, A. H., Wieskopf, J. S., Acland, E. L., Dokova, A., Kadoura, B., Leger, P., Mapplebeck, J. C. S., McPhail, M., Delaney, A., Wigerblad, G., Schumann, A. P., Quinn, T., Frasnelli, J., Svensson, C. I., … Mogil, J. S. (2014). Olfactory exposure to males, including men, causes stress and related analgesia in rodents. Nature Methods, 11(6), 629–632. https://doi.org/10.1038/nmeth.2935

Tashima, R., Koga, K., Sekine, M., Kanehisa, K., Kohro, Y., Tominaga, K., Matsushita, K., Tozaki-Saitoh, H., Fukazawa, Y., Inoue, K., Yawo, H., Furue, H., & Tsuda, M. (2018). Optogenetic Activation of Non-Nociceptive Aβ Fibers Induces Neuropathic Pain-Like Sensory and Emotional Behaviors after Nerve Injury in Rats. ENeuro, 5(1). https://doi.org/10.1523/ENEURO.0450-17.2018

Usoskin, D., Furlan, A., Islam, S., Abdo, H., Lönnerberg, P., Lou, D., Hjerling-Leffler, J., Haeggström, J., Kharchenko, O., Kharchenko, P. V, Linnarsson, S., & Ernfors, P. (2015). Unbiased classification of sensory neuron types by large-scale single-cell RNA sequencing. Nature Publishing Group, 18(1), 145–153. https://doi.org/10.1038/nn.3881

Warwick, C., Cassidy, C., Hachisuka, J., Wright, M. C., Baumbauer, K. M., Adelman, P. C., Lee, K. H., Smith, K. M., Sheahan, T. D., Ross, S. E., & Koerber, H. R. (2021). Mrgprd Cre lineage neurons mediate optogenetic allodynia through an emergent polysynaptic circuit. Pain, 162, 2120–2131. https://doi.org/10.1097/j.pain.0000000000002227

Weinreb, C., Abdal, M., Osman, M., Zhang, L., Lin, S., Pearl, J., Annapragada, S., Conlin, E., Gillis, W. F., Jay, M., Ye, S., Mathis, A., Mathis, M. W., Pereira, T., Linderman, S. W., & Datta, S. R. (2023). Keypoint-MoSeq: parsing behavior by linking point tracking to pose dynamics. BioRxiv, 2023.03.16.532307. https://doi.org/10.1101/2023.03.16.532307

Woolf, C. J. (2020). Capturing Novel Non-opioid Pain Targets. Biological Psychiatry, 87(1), 74–81. https://doi.org/10.1016/J.BIOPSYCH.2019.06.017

Xie, Y. F., Wang, J., & Bonin, R. P. (2018). Optogenetic exploration and modulation of pain processing. Experimental Neurology, 306, 117–121. https://doi.org/10.1016/J.EXPNEUROL.2018.05.003

Zhou, X., Wang, L., Hasegawa, H., Amin, P., Han, B. X., Kaneko, S., He, Y., & Wang, F. (2010). Deletion of PIK3C3/Vps34 in sensory neurons causes rapid neurodegeneration by disrupting the endosomal but not the autophagic pathway. Proceedings of the National Academy of Sciences of the United States of America, 107(20), 9424–9429. https://doi.org/10.1073/PNAS.0914725107

